# Highly contiguous assemblies of 101 drosophilid genomes

**DOI:** 10.1101/2020.12.14.422775

**Authors:** Bernard Y. Kim, Jeremy R. Wang, Danny E. Miller, Olga Barmina, Emily Delaney, Ammon Thompson, Aaron A. Comeault, David Peede, Emmanuel R. R. D’Agostino, Julianne Pelaez, Jessica M. Aguilar, Diler Haji, Teruyuki Matsunaga, Ellie E. Armstrong, Molly Zych, Yoshitaka Ogawa, Marina Stamenković-Radak, Mihailo Jelić, Marija Savić Veselinović, Marija Tanasković, Pavle Erić, Jian-jun Gao, Takehiro K. Katoh, Masanori J. Toda, Hideaki Watabe, Masayoshi Watada, Jeremy S. Davis, Leonie C. Moyle, Giulia Manoli, Enrico Bertolini, Vladimír Košťál, R. Scott Hawley, Aya Takahashi, Corbin D. Jones, Donald K. Price, Noah Whiteman, Artyom Kopp, Daniel R. Matute, Dmitri A. Petrov

## Abstract

Over 100 years of studies in *Drosophila melanogaster* and related species in the genus *Drosophila* have facilitated key discoveries in genetics, genomics, and evolution. While high-quality genome assemblies exist for several species in this group, they only encompass a small fraction of the genus. Recent advances in long read sequencing allow high quality genome assemblies for tens or even hundreds of species to be generated. Here, we utilize Oxford Nanopore sequencing to build an open community resource of high-quality assemblies for 101 lines of 95 drosophilid species encompassing 14 species groups and 35 sub-groups with an average contig N50 of 10.5 Mb and greater than 97% BUSCO completeness in 97/101 assemblies. These assemblies, along with detailed wet lab protocol and assembly pipelines, are released as a public resource and will serve as a starting point for addressing broad questions of genetics, ecology, and evolution within this key group.

## Introduction

The biological and genetic tractability of fruit flies (*Drosophila* and related genera) has led to their status as a premier model system for biological research, particularly in the genomic era (Clark et al., 2007; Hales et al., 2015). Current publicly available genome assemblies number in the tens of species, some with accompanying gene expression and regulation databases (Chen et al., 2014; modENCODE Consortium et al., 2010), comparative genomics tools (Stark et al., 2007), or population genomic data (Guirao-Rico & González, 2019; Lack et al., 2016; Signor et al., 2018). Unfortunately, these genomic resources are far from comprehensive for this remarkably biodiverse group, which encompasses over 1,600 described species (O’Grady & DeSalle, 2018). Expanding the phylogenetic scope of these resources will enable further study of the ecological and evolutionary forces that shape this large and diverse clade.

Recent developments in long-read sequencing make it increasingly feasible to quickly generate high-quality genomes at the level of whole clades. Long-read sequencing simplifies many genome assembly challenges by fully spanning complex regions, such as repetitive elements (generally <10kb in length), and allows generation of chromosome-level assemblies at a reasonable cost. A number of recent studies have used long-read technology to assemble high-quality *Drosophila* genomes for several species groups (Bracewell et al., 2019; Chakraborty et al., 2019; Hill et al., 2020; Mai et al., 2020; Miller et al., 2018). Notably, Miller et al. (2018) estimated the cost of a high-quality long-read assembly at US $1,000, a significant milestone in the democratization of large genome assembly projects.

Here we improve upon that benchmark to present a community resource of 101 *de novo* genome assemblies, from 95 drosophilid species contributed by *Drosophila* researchers from across the world, representing a diversity of ecologies and geographical distributions. We used a hybrid assembly approach with Oxford Nanopore (ONT) long-read sequencing to construct the draft genome and Illumina short reads for polishing. The quality of these genomes is assessed. We propose that under ideal conditions, at least two samples of a typical *Drosophila* genome can be sequenced per ONT release 9.4.1 (rev D) flow cell for as little as $350 (USD) per final high-quality genome. In conjunction with this manuscript and data, we provide a wet lab protocol on Protocols.io specifically optimized for *Drosophila* genome assembly, along with containerized computational pipelines on GitHub. These genome assemblies and technical resources should facilitate the process of conducting large-scale genome projects in this key model clade and beyond.

## Results & Discussion

### Taxon sampling

Briefly, our selection of species and strains for sequencing (**Table 1**) improves the geographic, ecological, and phylogenetic diversity of *Drosophila* genomes available to the public. The lines sequenced here represent 13 species groups (Toda, 2020) in both major subgenera (*Drosophila* and *Sophophora*); originate from mainland and island locations in North America, Europe, Africa, and Asia; are found from northern (e.g., *D. tristis, D*.*littoralis*) to equatorial (e.g., *D. bocqueti*) latitudes; represent notable independent transitions to herbivory (*Scaptomyza* and *Lordiphosa*); and include samples of a pest (*Zaprionus indianus*) taken from its native and invasive range. For some species, for instance some *Lordiphosa* spp., only wild-caught flies were sequenced. We sequenced lines in use for active research projects so additional resources like gene expression or population data are expected in the near future. Despite our efforts to improve species diversity, we acknowledge that this initial sampling is heavily biased towards taxa that can be maintained in the lab. Additional details of sample collection are provided in the **Methods**. Future work to improve biological and taxonomic diversity, particularly for species difficult to culture, should employ single fly sequencing and assembly workflows (Adams et al., 2020).

**Table 1.**
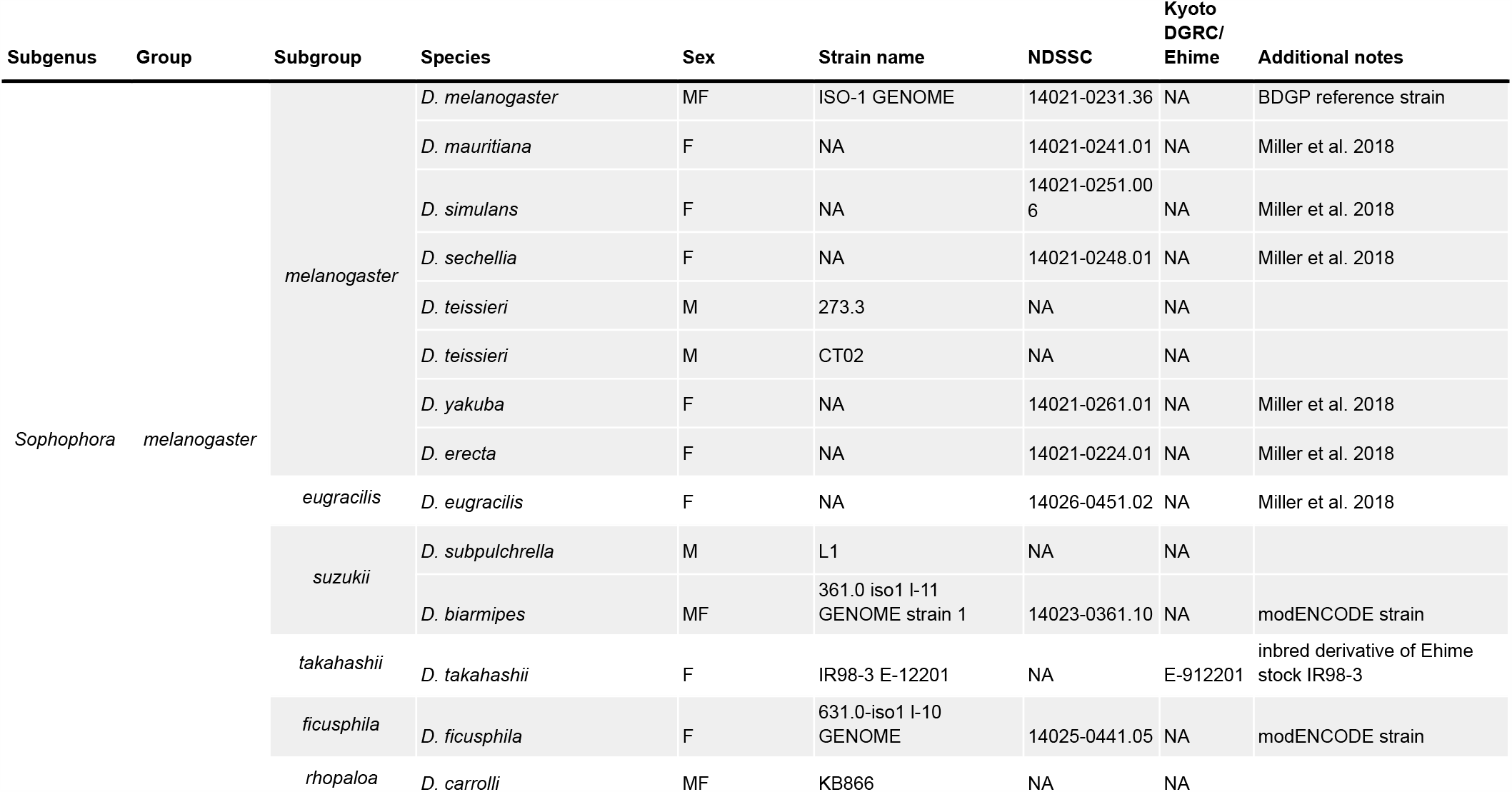
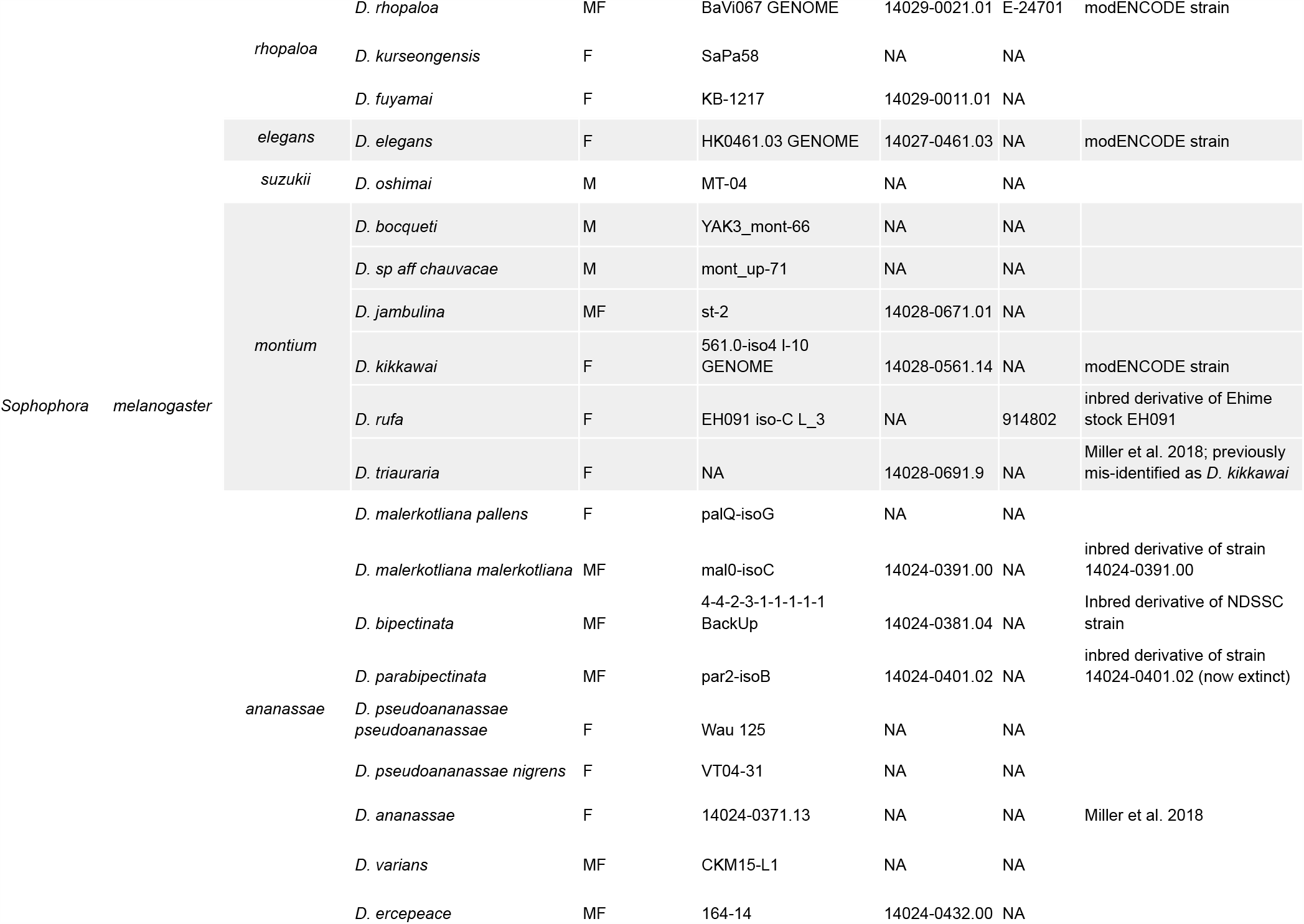
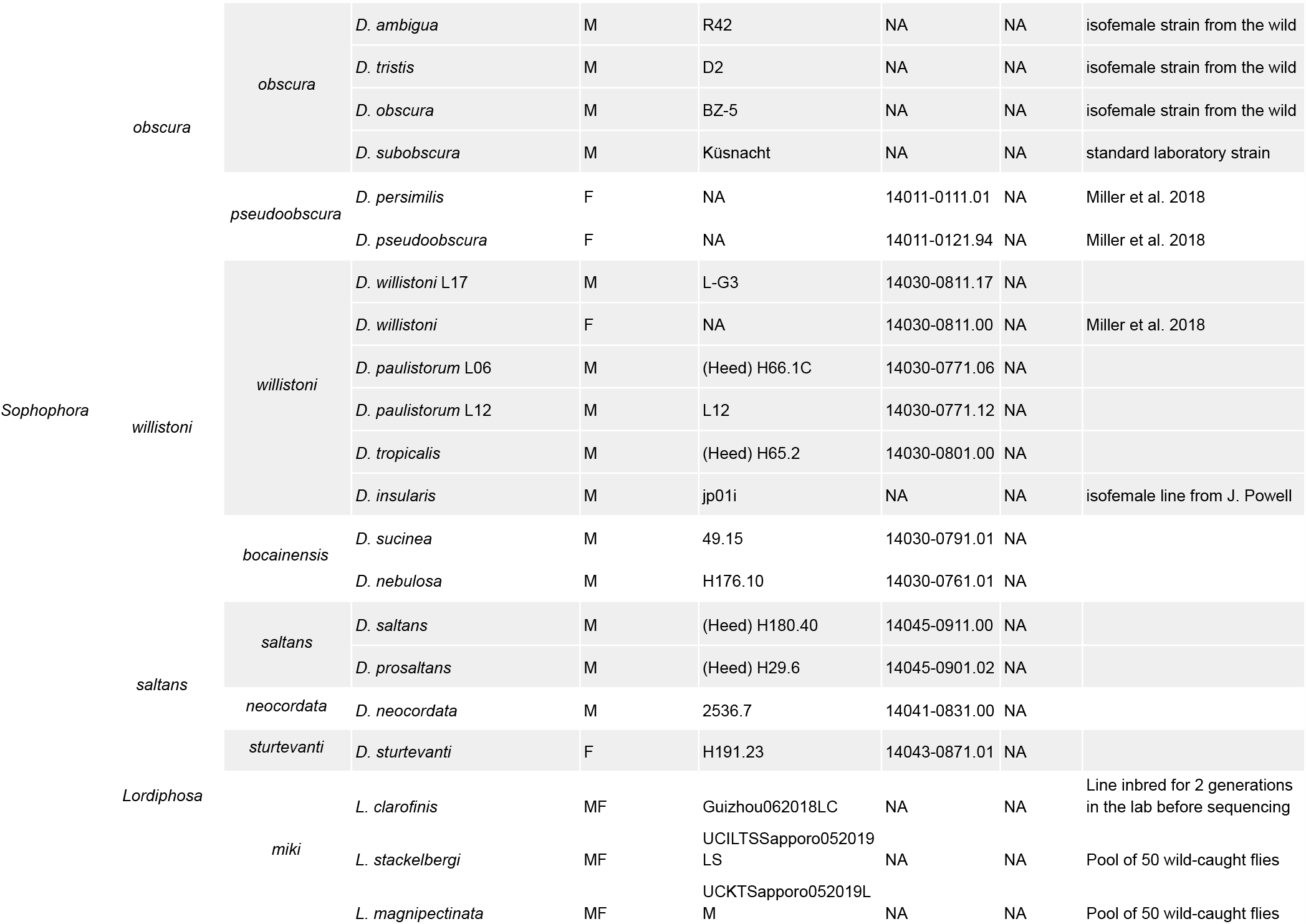
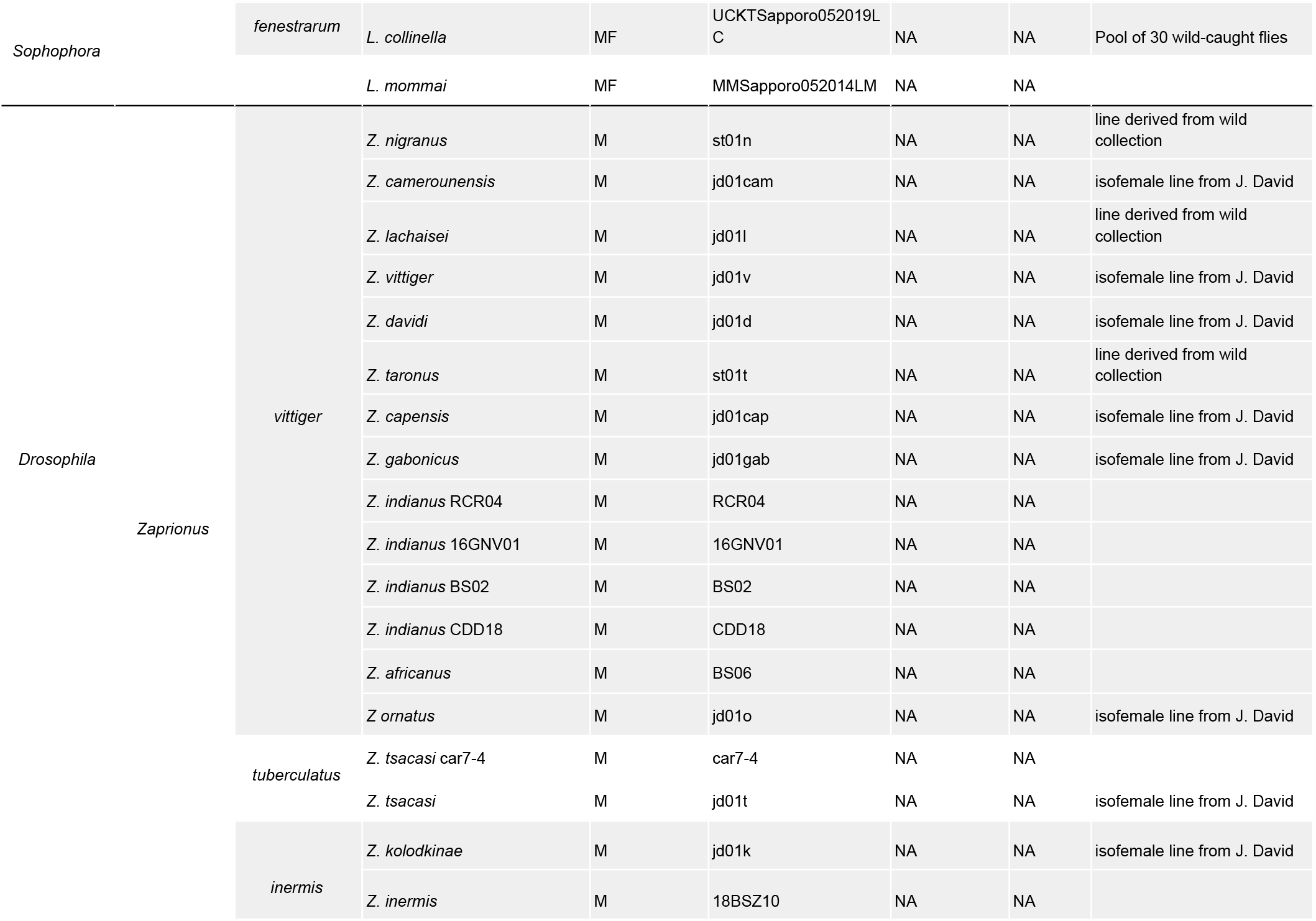
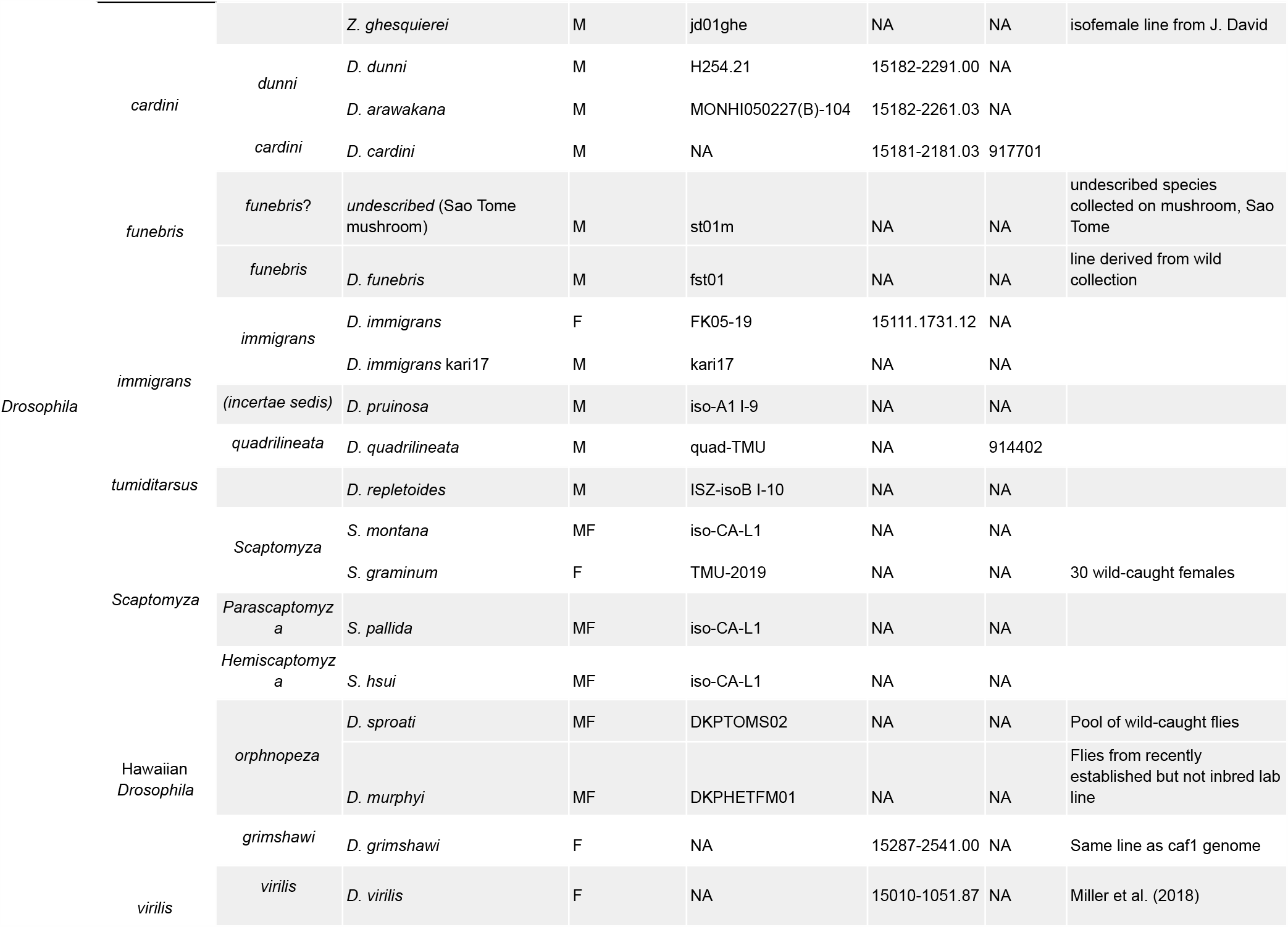
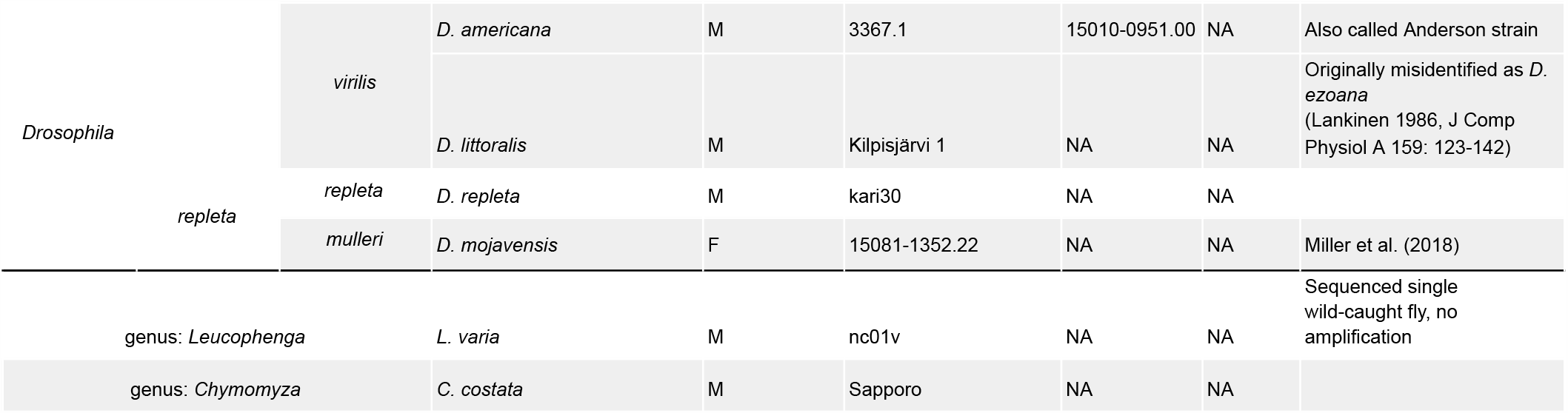
Species and strain information for all samples assembled for this work. Note: Species group and subgroup information is taken from the NCBI Taxonomy Browser with slight modifications following O’Grady and DeSalle (2018). Strain names along with corresponding NDSSC and Kyoto DGRC stock center numbers are provided to the best of our knowledge. See **Tables S1 and S3** for detailed information on samples and data.

### Near chromosome-scale assembly with ultra-long reads

We sequenced 101 fly strains using a modified ONT 1D ligation kit approach, optimized for DNA extractions from 15–30 whole flies and to reduce library prep cost while balancing read lengths and overall throughput. Sequencing runs varied with sample quality and type, and in general read lengths increased over the course of this work. Under optimal conditions, libraries prepared with the supplied protocol should yield 12–15 Gb of data per R9.4.1 flow cell with a read N50 greater than 20kb, and about 30% of data in reads longer than 50kb. We generated paired-end, 150bp Illumina reads for most strains unless public datasets were available.

Deep (average 52×) sequencing coverage with a substantial fraction of ultra-long (50-100 kb+) ONT reads (**Table S1**) resulted in highly contiguous genome assemblies (**Figure S1**) comparable to existing reference genomes in contiguity and completeness (**Figure 1, Table S2**). We used Flye (Kolmogorov et al., 2019) based on superior assembly contiguity and favorable runtimes relative to Miniasm (Li, 2016) and Canu (Koren et al., 2017) (**Figure S2, Methods**). Of 101 total assemblies, 94 contain over 98% of the assembly in contigs larger than 10kb, and contig N50s exceed 1 Mb for all but 7. In cases where DNA was extracted from pools of wild-caught flies or a single fly (*Leucophenga varia*) resulting in sub-optimal read lengths and output, the assembly was comparable to existing short read assemblies (**Figure 1A & 1B**). High contiguity resulted in BUSCO completeness in the range of 97–99+% for all but the 4 most fragmented genomes (**Figure 1C**).

**Figure 1.**
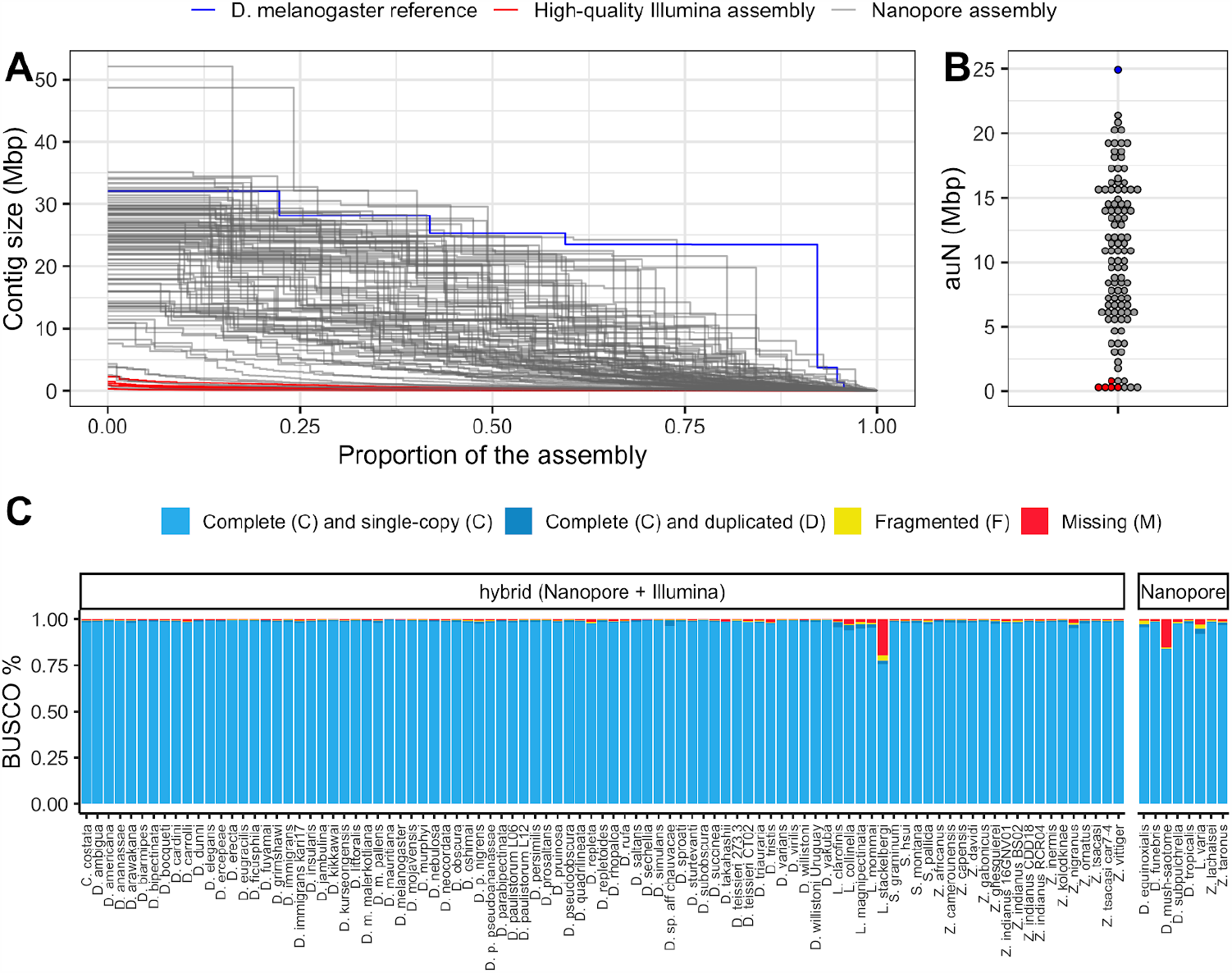
Nanopore-based assemblies are highly contiguous and complete. (**A,B**) Assembly contiguity is compared to the *D. melanogaster* v6.22 reference genome (blue) as well as 5 recently published, highly contiguous Illumina assemblies (red lines, *D. birchii, D. bocki, D. bunnanda, D. kanapiae, D. truncata*; Bronski et al., 2020). (**A**) *Nx* curves, or the (y-axis) size of each contig when contigs are sorted in descending size order, in relation to the (x-axis) cumulative proportion of the genome assembly that is covered. (**B**) The distribution of the *auN* statistic, a measure of contiguity obtained by calculating the area under the *Nx* curve. A contig break will always result in lower *auN* but not necessarily a lower *N50*. (**C**) Assembly completeness assessed by BUSCO v4.0.6 (Simão et al., 2015). Note, *D. equinoxialis* was evaluated with BUSCO v3.0.2 due to an unfixable bug. Individual assembly summary statistics are provided in **Table S2**.

While we estimate the sequencing cost of a single genome assembly, under the typical conditions presented here, to be $350 (USD), there are opportunities for further optimization in future work. Currently, sequencing runs optimized for ultra-long reads suffer from low throughput due to pore clogging during sequencing. We find that near chromosome level contiguity can be achieved even with minimal (∼10×) coverage of reads longer than 25kb (**Figure S3**). Additional read depth will improve consensus sequence accuracy, important for downstream tasks like annotation. However this can be obtained from shorter Nanopore and Illumina reads, which are both easier and cheaper to generate.

### A comparative genomics resource

To demonstrate the potential this dataset holds for the study of genome evolution and chromosome organization, we revisit a classic result with our highly contiguous assemblies. Although the ordering of genes in drosophilid chromosomal (Muller) elements has been extensively shuffled throughout ∼53 million years of evolution (Suvorov et al., *in prep*), the gene content of each element remains largely conserved (Bracewell et al., 2019; Ranz et al., 2001; Sturtevant & Novitski, 1941). To examine synteny in our assemblies, many of which contain several contigs tens of megabases in length, we constructed an undirected graph using single-copy orthologous markers (i.e., BUSCOs). The number of times two markers were connected by assemblies determined the weight of the graph’s edges. When a graph layout method was applied to visualize these relationships (**Methods**), we found that orthologs clustered by the *D. melanogaster* chromosome on which they are found, consistent with the expected conservation of gene content in Muller elements across drosophilids. Furthermore, the lack of clear order within groups is consistent with extensive shuffling within Muller elements. This demonstrates that our dataset can be used for studies of genome evolution. New reference-free, whole-genome alignment methods (Armstrong et al., 2020) should substantially facilitate these kinds of comparative analyses.

### Repeat content

A large number of genome assemblies enables comparative analysis of repeat variation against a wide range of genome sizes (140–450Mb), for example the independent expansions of satellite repeats in *D. grimshawi* or retroelements in *D. paulistorum, D. bipectinata*, or *D. subpulchrella* (**Figure 3**). Within our dataset alone, RepeatMasker annotations show large variation in repeat content among drosophilids (**Figure 3**). No correlation exists between assembly contiguity and repeat content (**Figure S4**), suggesting long-read sequencing overcomes many of the challenges to drosophilid genome assembly posed by repetitive sequences. Additionally, we observe a positive relationship between the size of repetitive sequences and non-repetitive sequences, suggesting that genome size is influenced by expansions and contractions of both portions of the genome (**Figure S5**). The high continuity of these assemblies should allow the identification of complete transposable elements in the genomes and allow for the analyses of transposable element evolution at the level of individual transposable elements or transposable element families in a way that is not feasible with more fragmented genome assemblies (Clark et al., 2007).

**Figure 2.**
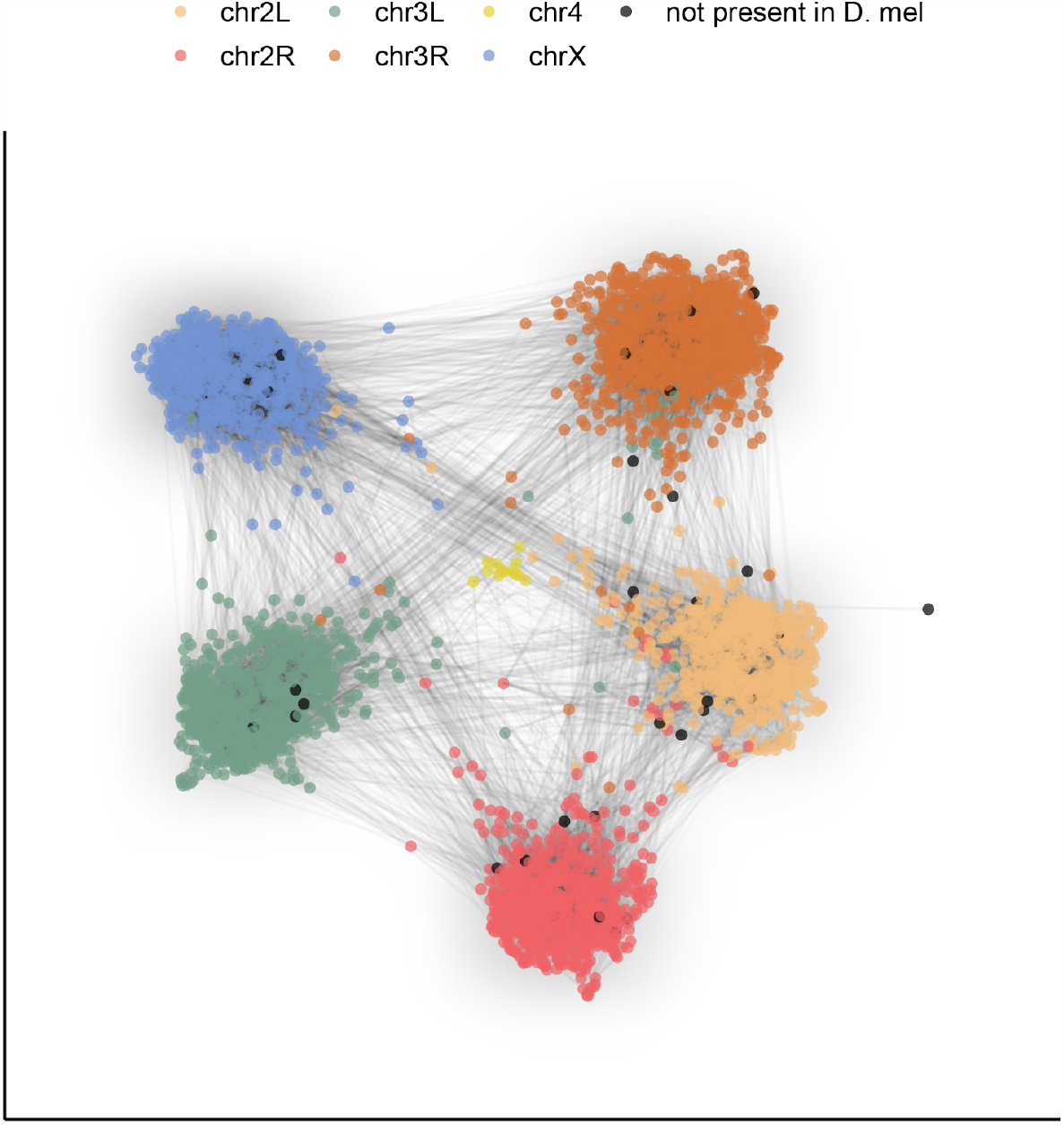
Gene content of Muller elements is conserved across drosophilids while gene order changes. Each node in this graph represents an orthologous marker corresponding to single-copy orthologs annotated by BUSCOv4 (Seppey et al., 2019; Simão et al., 2015). An edge between two nodes represents the number of times that BUSCO pair is directly connected within an assembly. Each BUSCO is colored by the chromosome arm in *D. melanogaster* that it is found on. The ForceAtlas2 (Jacomy et al., 2014) graph layout algorithm was used for visualization.

**Figure 3.**
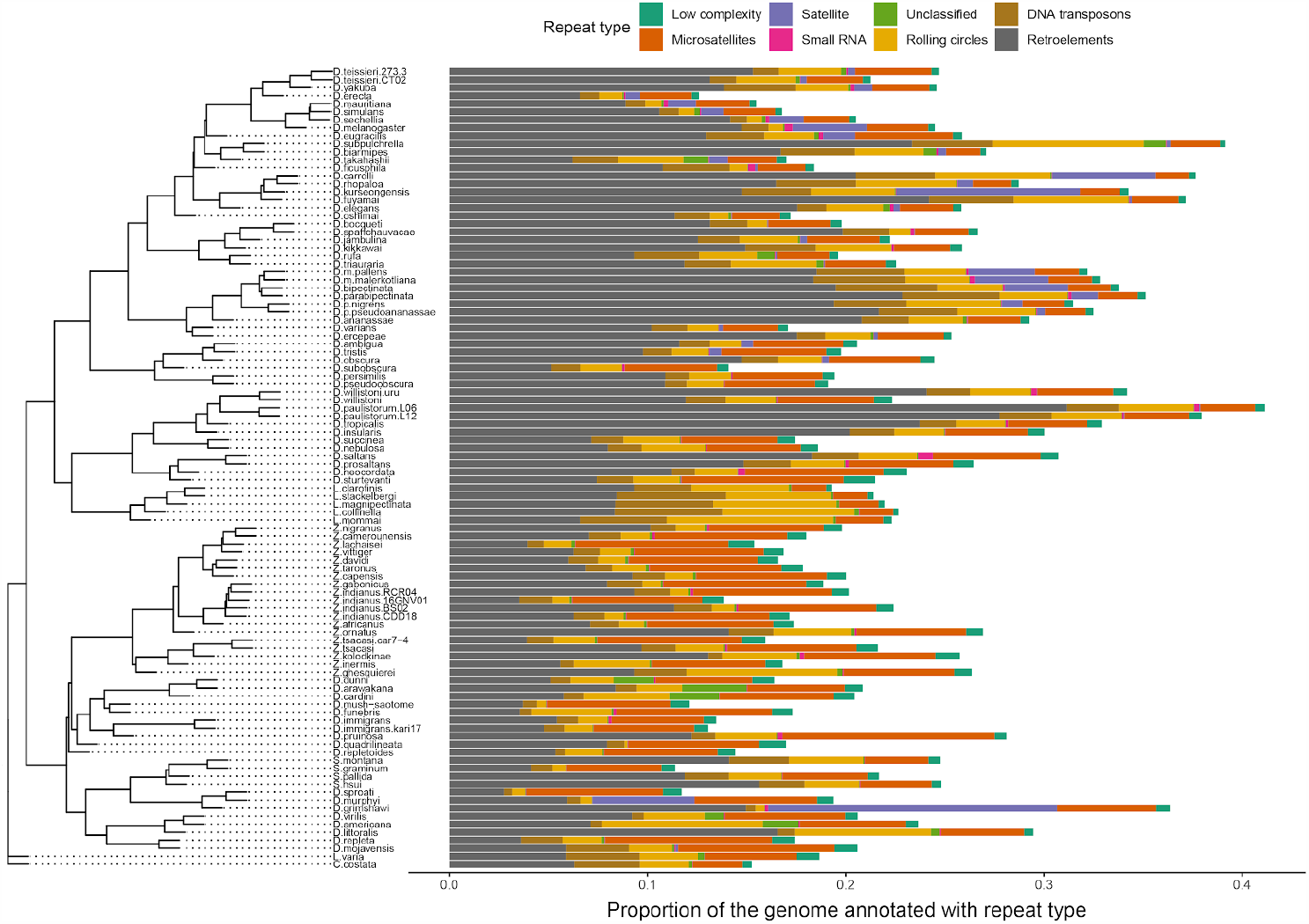
Repeat content varies greatly between drosophilid groups. For each species, the proportion of each genome annotated with a particular repeat type is depicted. Species relationships were inferred by randomly selecting 250 of the set of BUSCOs (Simão et al., 2015) that were complete and single-copy in all assemblies. RAxML-NG (Kozlov et al., 2019) was used to build gene trees for each BUSCO then ASTRAL-MP (Yin et al., 2019) to infer a species tree. Repeat annotation was performed with RepeatMasker (Smit et al., 2013) using the Dfam 3.1 (Hubley et al., 2016) and RepBase RepeatMasker edition (Bao et al., 2015) databases.

### Reproducibility

Detailed laboratory protocols, computational pipelines, and computational container recipes are provided as a reference and to maximize reproducibility. The protocol is publicly available at Protocols.io (dx.doi.org/10.17504/protocols.io.bdfqi3mw) and pipeline scripts along with associated compute containers are provided in a public GitHub repository (https://github.com/flyseq/drosophila_assembly_pipelines). See **Methods** for additional details on compute containers.

### Future directions

We have described an open community resource of 101 nearly chromosome-level drosophilid genome assemblies, adding to or improving upon many assemblies already available for this group (Suvorov et al., n.d.) as well as providing detailed protocols for adding additional genomes economically and easily. We envision the provided dataset being used to address a large number of outstanding questions entailing large comparative analyses among species and to improve the genome annotation process for these species (Armstrong et al., 2020; Fiddes et al., 2018; Shumate & Salzberg, 2020). Finally, these data will allow population genomic data to be compared for a large number of species, providing unprecedented resolution to investigate fundamental questions about the evolutionary process.

## Materials and Methods

### Taxon Sampling and Sample Collection

The selection of species used for this study was driven by several key objectives. First, we aimed to provide data for ongoing research projects. Second, we aimed to supplement existing genomic data, both as a benchmarking resource against well-studied references (e.g. *D. melanogaster*) and to provide a technological update to some older assemblies (modENCODE Consortium et al., 2010). Third, we aimed to increase the phylogenetic and ecological diversity of publically available *Drosophila* genome assemblies.

In most cases, genomic DNA was collected from lab-raised flies, which were either derived from lines maintained at public *Drosophila* stock centers and individual labs or, in a few cases, from F1 or F2 progeny of flies recently collected in the wild. We collected specimens from the wild with standard fruit or mushroom-baited traps, sweep netting, and aspiration. We established isofemale lines from individual females collected using these baits unless otherwise specified (**Table S1**). For species difficult to culture in the lab (all *Lordiphosa* spp. except *Lo. clarofinis, D. sproati, D. murphyi, Le. varia, S. graminum*), either wild-caught flies or flies from a transient lab culture were used. In accordance with domestic and international shipping laws, these flies were either fixed in ethanol before transport (*Lordiphosa* spp., *D. subobscura, D. obscura, C. costata, D. littoralis, D. tristis, D. ambigua*) or transported with permits (P526P-15-02964 to D. Matute, P526P-20-02787 and P526P-19-01521 to A. Kopp, and Hawaii State permit I1302 to D. Price).

Of 101 total assemblies, we include 13 genomes assembled with re-analyzed sequences from Miller *et al*. (Miller et al., 2018); 60 genomes from stock center lines or established lab cultures; 22 genomes from lab-raised flies derived from recent wild collections; and 6 genomes from wild-caught flies. Of note, 6 *Zaprionus* lines used in this study (*Z. africanus, Z. indianus, Z. tsacasi, Z. nigranus, Z. taronus*) were assembled by Comeault et al. (Comeault et al., 2020), but updated higher contiguity assemblies are provided with this manuscript with the exception of *Z. indianus* line 16GNV01 (see “Alternative hybrid assembly process” section below). Details on each sample including (if available) line designations and collection information, are provided in **Table S3**.

### DNA extraction and Nanopore sequencing

A high molecular weight (HMW) genomic DNA (gDNA) extraction and Nanopore library prep was performed for each sample, with slight variation in the protocol through time and to deal with differences in sample quality or preservation. Here, we briefly describe a recommended general protocol for HMW gDNA extraction and library prep from 15–30 flies. This protocol is sufficient to reproduce all results from this manuscript at the same or higher levels of data quality. Detailed step-by-step instructions are provided at Protocols.io (dx.doi.org/10.17504/protocols.io.bdfqi3mw). We note one exception made necessary by sample availability and shipping laws. *Scaptomyza graminum* gDNA was extracted by using the Qiagen Blood & Cell Culture DNA Mini Kit from 30 unfrozen flies and prepared with the ONT LSK109 kit without any modifications to the manufacturer’s instructions.

Genomic DNA was prepared from about 30 flash frozen or ethanol fixed adult flies. For non-inbred samples, we tried to use 15 flies or less to minimize the genetic diversity of the sample. In the absence of amplification, about 1.5–3 μg of input DNA is needed to prepare 3–4 library loads with the ONT LSK109 kit. Sufficient input DNA is particularly important when selecting for longer reads. Ethanol preserved samples were soaked in a rehydration buffer (400 mM NaCl, 20 mM Tris-HCl pH 8.0, 30 mM EDTA) for 30 minutes at room temperature (∼23°C), dabbed dry with a Kimwipe, then frozen for 1 hour at -80°C before extraction. Frozen flies were ground in 1.5 mL of homogenization buffer (0.1M NaCl, 30mM Tris HCl pH 8.0, 10 mM EDTA, 0.5% Triton X-100) with a 2 mL Kontes Dounce homogenizer. The homogenate was centrifuged for 5 minutes at 2,000 ×*g*, the supernatant discarded by decanting, and the pellet resuspended in 100 μL of fresh homogenization buffer. This mixture was then added to a tube with 380 μL extraction buffer (0.1M Tris-HCl pH 8.0, 0.1M NaCl, 20mM EDTA) along with 10 μL Proteinase K (20 mg/mL), 10 μL SDS (10% w/v), and 2 μL RNAse A (10 mg/mL). This tube was incubated at 50°C for 4 hours, with mixing at 30–60 minute intervals by gentle inversion.

High molecular weight gDNA was purified with a standard phenol-chloroform extraction. The lysate was extracted twice with an equal volume of phenol chloroform isoamyl alcohol (25:24:1 v/v) in a 2 mL light phase lock gel tube. Next, the aqueous layer was decanted into a fresh 2mL phase lock gel tube then extracted once with an equal volume of chloroform. The use of the phase lock gel tube reduces DNA shearing at this stage by minimizing pipette handling. HMW DNA was precipitated by adding 0.1 volume of 3M sodium acetate and 2.0–2.4 volumes of cold absolute ethanol. Gentle mixing resulted in the precipitation of a white, stringy clump of DNA, which was then transferred to a DNA LoBind tube and washed twice with 70% ethanol. After washing, the DNA was pelleted by centrifugation and all excess liquid removed from the tube. The pellet was allowed to air dry until the moment it became translucent, resuspended in 65 μL of 1× Tris-EDTA buffer on a heat block at 50°C for 60 minutes, then incubated for at least 48 hours at 4°C. After 48 hours, the viscous DNA solution was mixed by gentle pipetting with a P1000 tip. This controlled shearing step encourages resuspension of HMW DNA and improves library prep yield. DNA was quantified with Qubit and Nanodrop absorption ratios were checked to ensure 260/280 was greater than 1.8 and 260/230 was greater than 2.0.

The sequencing library was prepared following the ONT Ligation Sequencing Kit (SQK-LSK109) protocol, with two important modifications. First, we started with approximately 3 μg of input DNA, three times the amount recommended by the manufacturer. Second, we utilized size-selective polymer precipitation (Paithankar & Prasad, 1991) with the Circulomics Short Read Eliminator (SRE) buffer plus centrifugation to isolate DNA instead of magnetic beads. We found this to be necessary because magnetic beads irreversibly clumped with viscous HMW gDNA, decreasing library yield and limiting read lengths. The manner in which this was performed was specific to the cleanup step. After the end-prep/repair step, the SRE buffer was used according to the manufacturer’s instructions. After adapter ligation, DNA was pelleted by centrifuging the sample at 10,000×*g* for 30 minutes without the addition of any reagents, since DNA readily precipitated upon addition of the ligation buffer. Ethanol washes were avoided past this step since ethanol will denature motor proteins in the prepared library. Instead, the DNA pellet was washed with 100 μL SFB or LFB (interchangeably) from the ligation sequencing kit instead of 70% ethanol. If library yield was sufficient (>50 ng/μL), the Circulomics SRE buffer was used for a final round of size selection, replacing the ethanol wash with LFB/SFB as described above. Of note, a cheaper and open-source alternative made with polyethylene glycol MW 8000 (PEG 8000), although less effective at size selection, to the SRE buffer is described by Tyson (Tyson, 2020) (dx.doi.org/10.17504/protocols.io.7euhjew). A 1:1 dilution of the PEG 8000 solution described in that protocol can be substituted for SFB or LFB in the washing steps described above.

The typical yield of a library prepared in this manner is in the range of 1–1.5 μg. Approximately 350 ng of the prepared library was loaded for each sequencing run. To maintain flow cell throughput and read length, flow cells were flushed every 8–16 hours with the ONT Flow Cell Wash Kit (EXP-WSH003) and reloaded with a fresh library.

### Short read data for polishing

We performed 2×150 bp Illumina sequencing for most of the strains that did not have publicly available short read data available. Illumina libraries were prepared from the same gDNA extractions as the Nanopore library for most samples, with some exceptions as described in **Table S2**. The libraries were prepared in either of two manners. For the majority of samples, sequencing libraries were prepared with a modified version of the Nextera DNA Library Kit protocol (Baym et al., 2015) and sequencing was performed by Admera Health on NextSeq 4000 or HiSeq 4000 machines. Alternatively, Illumina libraries were prepared with the KAPA Hyper DNA kit according to the manufacturer’s protocol and sequenced at the UNC sequencing core on a HiSeq 4000 machine. In either case, all samples on a lane were uniquely dual indexed. Illumina sequencing was not performed for *D. equinoxialis, D. funebris, D. subpulchrella, D. tropicalis, Le. varia, Z. lachaisei, Z. taronus*, and the unidentified São Tomé mushroom feeder due to material unavailability (line extinction/culling). Details for each sample, including accession numbers for any public data used in this work, are provided in **Table S2**.

### Choice of long read assembly program

Flye v2.6 (Kolmogorov et al., 2019) was used due to its quick CPU runtime, low memory requirements, excellent assembly contiguity, and its consistent performance on benchmarking datasets (Wick & Holt, 2020). We additionally validated the performance of Flye for *Drosophila* genomes using Nanopore data previously generated by Miller *et al*. (2018) and 60× depth of new Nanopore sequencing of the Berkeley Drosophila Genome Project ISO-1 strain of *D. melanogaster*. We assembled genomes with Flye v2.6 and Canu v1.8 (Koren et al., 2017) to evaluate simple benchmarks of assembly contiguity and run time and to provide a comparison to the Miniasm (Li, 2016) assemblies from Miller *et al*. (Miller et al., 2018) Canu produced relatively contiguous assemblies, but a single assembly took several days on a 92-core cloud server and even longer when a large number of extra-long (>50kb) reads were present in the data. This was determined to be too costly when scaled to >100 species. In addition to a much shorter (8–12 hours wall-clock time) runtime, Flye also produced significantly more contiguous assemblies than those reported by Miller *et al*. (**Figure S2**). Note, several new long read assemblers have been released and these assembly programs have been significantly updated since this work was performed. Assembler performance should be evaluated with up-to-date versions in any future work.

### Assembly and long read polishing

After Nanopore sequencing was performed, raw Nanopore data were basecalled with Guppy v3.2.4, using the high-accuracy caller (option: -c dna_r0.4.1_450bps_hac.cfg). Raw Nanopore data previously generated by Miller *et al*. (Miller et al., 2018) were processed in the same manner.

Next, basecalled reads were assembled using Flye v2.6 with default settings. Genome size estimates (option: --genomeSize) were obtained through a web search or taken from a closely related species. If no such information was available, an initial estimate of 200 Mb was used. The specific genome size estimate is provided in **Table S2**.

After generating a draft assembly, we performed long read polishing using Medaka (https://nanoporetech.github.io/medaka/draft_origin.html). Reads were aligned to the draft genome with Minimap2 v2.17 (Li, 2016) before each round of polishing (option: -ax ont). The draft was polished with two rounds of Racon v1.4.3 (Vaser et al., 2017) (options: -m8 -x 6 -g 8 -w 500) and then a single round of Medaka v0.9.1.

### Haplotig identification and removal

Next, we assessed each assembly for the presence of multiple haplotypes (haplotigs) using BUSCO v3.0.2 (Simão et al., 2015; Waterhouse et al., 2018) on the Medaka-polished sequences. If the BUSCO duplication rate exceeded 1%, haplotig identification and removal was performed, but on the draft assembly produced by Flye rather than the polished assembly. Purge_haplotigs v1.1.1 (Roach et al., 2018) was run on these sequences following the guidelines provided by the developer (https://bitbucket.org/mroachawri/purge_haplotigs). Illumina reads were mapped to the draft assembly with Minimap2 (option: -ax sr) to obtain read depth information. The optional clipping step was performed to remove overlapping (duplicate) contig ends. Finally, remaining contigs were re-scaffolded with Nanopore reads using npScarf v1.9-2b (Cao et al., 2017), with support from at least 4 long reads required to link two contigs (option: --support=4). These sequences were polished with Racon and Medaka as described above.

### Final polishing and decontamination

The Medaka-polished assembly was further polished with Illumina data and any contigs identified as microbial sequences were removed. For each polishing round, Illumina reads were mapped to the draft assembly with Minimap2 (option: -ax sr) and polished with Pilon v1.23 (Walker et al., 2014). If a genome did not have an accompanying short read dataset but Illumina reads were available from a different strain of the same species (**Table S1**), Pilon was run without correcting SNVs (option: --fix indels,gaps,local). After Pilon polishing, assembly completeness was assessed again with BUSCO v3.0.2. We used BLAST (version 2.10.0) (Altschul et al., 1990) to remove any contigs not associated with at least one BUSCO that were also of bacterial, protozoan, or fungal origin. Finally, any sequences flagged by the NCBI Contamination Screen were excluded or trimmed.

### Alternative hybrid assembly process

*Zaprionus indianus* line 16GNV01 had insufficient Nanopore data for a Flye assembly. For this line only and to consolidate all assemblies as a single resource, the same genome assembly from Comeault *et al*. (Comeault et al., 2020) is both reported here and associated with the NCBI BioProject associated with this work. An alternative assembly strategy was taken for this line (Comeault et al., 2020). Briefly, short-read sequence data was assembled first using SPAdes v3.11.1 (Bankevich et al., 2012) using default parameters. Nanopore reads were corrected with Illumina data using FMLRC v.1.0.0 (Wang et al., 2018) and subsequently used to scaffold the SPAdes assembly using LINKS v.1.8.7 (Warren et al., 2015) using the recommended iterative approach of 33 iterations with incrementally increasing *k*-mer distance threshold. The resulting scaffolds were polished with four rounds of Racon followed by four rounds of Pilon (but without Medaka) as described above.

### Repeat annotation and masking

Each draft assembly was soft repeat masked with RepeatMasker v4.1.0 (Smit et al., 2013) at medium sensitivity, with both Dfam 3.1 (Hubley et al., 2016) and RepBase RepeatMasker edition (Bao et al., 2015) repeat libraries installed (options: --species drosophila --xsmall). RepeatMasker was initialized with cross_match v1.090518 (Green, 2009) as the sequence search engine and Tandem Repeat Finder v4.0.9 (Benson, 1999).

### Assessing assembly contiguity and completeness

Assembly contiguity statistics were computed using a series of custom shell and R scripts. Fasta files were parsed with Bioawk v1.0 and summary statistics were computed in the standard manner with custom scripts. Of particular note, we preferentially presented the *auN* statistic over contig *N50* to summarize assembly contiguity, although both are provided. The *auN* statistic is computed as the area under an *Nx* curve:

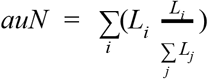

The advantage of *auN* for comparing assemblies is its sensitivity to assembly breaks: a break may not always affect *N50* but will always affect *auN* (Li, 2020).

Assembly completeness was assessed with BUSCO v4.0.6 (Seppey et al., 2019; Simão et al., 2015), in genome mode with the diptera_odb10 database (options: --m geno -l diptera_odb10 --augustus_species fly). Note that this BUSCO version is different (v4 vs v3). BUSCO v4 was released while the assemblies were in progress, and we wished to evaluate final completeness with the most up-to-date tools while retaining consistency across the assembly pipeline. BUSCO v4 (tested with v4.0.6-v4.1.3) runs with the *D. equinoxialis* genome were unsuccessful. For *D. equinoxialis* only, BUSCO v3.0.4 was run with the diptera_odb9 database (options: --m geno -l diptera_odb9).

### Species tree inference from BUSCO orthologs

We inferred species relationships using complete and single-copy orthologs identified by the BUSCO analysis. Amino acid sequences were used instead of nucleotide sequences to achieve better alignments in the face of high sequence divergence (Bininda-Emonds, 2005). Out of 990 single-copy orthologs present in all assemblies, we randomly selected 250 to construct gene trees. The predicted protein sequence of each ortholog was aligned separately with MAFFT v7.453 (Katoh & Standley, 2013), using the E-INS-i algorithm (options: --ep 0 --genafpair --maxiterate 1000). Gene trees were inferred with RAxML-NG v0.9.0 (Kozlov et al., 2019), using the Le and Gascuel (2008) amino acid substitution model (options: --msa-format FASTA --data-type AA --model LG). The summary method ASTRAL-MP v.5.14.7 (Yin et al., 2019) was run with default settings to reconstruct the species tree. We note that this is not intended to be a definitive phylogenetic reconstruction of species relationships; see Suvorov et al. (Suvorov et al., *in prep*) for a time-calibrated phylogeny utilizing 158 drosophilid whole genomes.

### Analysis of chromosome organization

Syntenic comparisons were performed by representing the genome assemblies as paths through an undirected graph. The path each genome traverses can be considered a series of connections between single copy orthologous markers (i.e., BUSCOs). Using BUSCO v4 annotations for each final genome, we constructed a 3,285 by 3,285 symmetric adjacency matrix, with row and column headers (nodes) corresponding to 3,285 possible BUSCOs from the diptera_odb10 database. Off-diagonal entries in each matrix (edges) were the number of times two single-copy BUSCOs were found as connected and immediate neighbors in the assemblies. Sequences of three or more BUSCOs were not considered. The graph was then visualized in two dimensions using the ForceAtlas2 graph layout algorithm (Jacomy et al., 2014) as implemented in the ForceAtlas2 R package (https://github.com/analyxcompany/ForceAtlas2). While this method is primarily designed for flexible, user-friendly tuning of graph visualization, it is similar in effect to other nonlinear dimensionality reduction techniques (Böhm et al., 2020). ForceAtlas2 was run with the settings: tolerance=1, gravity=1, iterations=3000. *D. equinoxialis* was omitted from this analysis due to the BUSCO v4 issues mentioned previously.

### Repeat content and genome size analysis

The contribution of repeat content to genome size variation in *Drosophila* was examined by comparing the number of bases in each genome annotated as a type of repeat (previously described) to the number of bases not annotated as repetitive sequence. Phylogenetic independent contrasts (Felsenstein, 1985) were computed for the counts of bases in both categories using the R package ape v5.4.1 (Paradis & Schliep, 2019) using the species tree described above with the root age set to 53 million years following the estimate in Suvorov et al. (Suvorov et al., n.d.).

### Compute containers

While the overall computational demands of this work were high, the unique computational challenge we faced was the variety of computational resources used for various stages of the assembly process. Assemblies took place across local servers, institutional clusters, and cloud computing resources. A key factor in ensuring reproducibility across computing environments was the use of computing containers, which is like a lightweight virtual machine that can be customized such that sets of programs and their dependencies are packaged together. Specifically, we used the programs Docker and Singularity to manage containers. These programs allow containers to be built and packaged as an image file which is transferred to another computer. A Dockerfile, a text file containing instructions to set up an image, is used to select the Linux operating system and the suite of programs to be installed within a Docker container. Singularity is used to package the Docker container as an image file that can be transferred to and used in a cluster or cloud environment without the need for administrative permissions. Standard commands are then run inside the container environment. The files and instructions necessary to build these containers, which will allow for the exact reproduction of the computing environment in which this work was performed, are provided at: https://github.com/flyseq/drosophila_assembly_pipelines. We hope these files will facilitate the work of researchers new to Nanopore sequencing or the genome assembly process.

## Data availability

Supplementary File 1: Figure S1 Flow chart depiction of the assembly pipeline. Supplementary File 2: Figure S2 Large improvements in assembly contiguity from an updated assembly workflow.

Supplementary File 3: Figure S3 Highly contiguous assemblies can be obtained with lower coverage of ultra-long reads.

Supplementary File 4: Assembly contiguity is not determined by repeat content. Supplementary File 5: The non-repetitive and repetitive portions of the genome both contribute to genome size differences in Drosophila.

Supplementary File 6: Table S1 Description of data used for this project, including accession numbers for public data.

Supplementary File 7: Table S2 Assembly summary statistics.

Supplementary File 8: Table S3 Detailed sample information.

Supplementary Files 1-8 are available at: https://doi.org/10.6084/m9.figshare.13377179

Whole genome sequencing data generated by this work are available at NCBI [BioProjectPRJNA675888](https://www.ncbi.nlm.nih.gov/bioproject/675888). Preliminary access to genome assemblies is provided at https://web.stanford.edu/~bkim331/files/genomes/. Raw Nanopore data are available by request.

Details are also provided at: http://flyseq.org/

Genome assembly pipeline and code: https://github.com/flyseq/drosophila_assembly_pipelines

Full laboratory protocol: dx.doi.org/10.17504/protocols.io.bdfqi3mw

## Acknowledgments

We thank Brandon Cooper, Antonio Serrato-Capuchina, and David Turissini for help with collections and field logistics; Sarah C.R. Elgin, Wilson Leung, Elena Gracheva, and Sophia Bieser for help with modENCODE fly lines; Jonathan Chang for helpful discussions about phylogenetic methods; Charlotte Helfrich-Förster for providing lab resources for G. Manoli and E. Bertolini; and John Tyson plus the staff at Circulomics, in particular Kelvin Liu and Michelle Kim, for many helpful discussions about long read library prep and sequencing.

